# Caspofungin resistance in *Candia auris* due to mutations in *Fks*1 with adjunctive role of chitin and key cell wall stress response pathway genes

**DOI:** 10.1101/2020.07.09.196600

**Authors:** Dipti Sharma, Raees A. Paul, Arunaloke Chakrabarti, Sanjay Bhattacharya, Rajeev Soman, Shamanth A Shankarnarayan, Dipali Chavan, Shreya Singh, Parijit Das, Harsimran Kaur, Anup K Ghosh, Shivaprakash M Rudramurthy

**Affiliations:** Department of Medical Microbiology, PGIMER, Chandigarh, India; Department of Microbiology, Tata Medical Centre, Kolkata, India; Jupiter Hospital, Pune, India; Tata Medical Centre, Kolkata, India

**Author notes:** **Corresponding author**: Shivaprakash M Rudramurthy, Professor, Mycology Division, Department of Medical Microbiology, Postgraduate Institute of Medical Education and Research, Chandigarh, India-160012, /2755155. **Co-corresponding author**: Arunaloke Chakrabarti, Professor, Mycology Division, Department of Medical Microbiology, Postgraduate Institute of Medical Education and Research, Chandigarh, India-160012, /2755155.

## Abstract

The emergence of echinocandin resistance in *C. auris* has become a major concern. Point mutations in Fks1 subunit of β-D-glucan synthase is the primary mechanism in echinocandin resistance. However, resistant isolates with wild type *Fks*1 are not infrequent. We screened 199 clinical *C. auris* isolates from 30 centres across India for echinocandin resistance. The cohort also contained six sequential isolates from a liver transplant recipient. Eleven isolates (5.7%) from 11 patients, and those six serial isolates had elevated echinocandin minimum inhibitory concentrations (MIC). Three of these 17 isolates carried S639F mutation in hot spot 1 region of *Fks1*. A novel *Fks*1 mutation, F635Y was identified in two resistant isolates, and a related F635L mutation was detected in four of the six sequential isolates. Resistant isolates (MIC≥2 mg/L) and those with intermediate caspofungin susceptibility (MIC, 1.0 mg/L) demonstrated higher induction of chitin synthase gene, *Chs*1 [resistant, 2.2(1.3-5.8); intermediate, 6(2.5-11.2)] compared to susceptible isolates [1.2(0.8-2), P<0.05]. However, the expression of the *Fks1* subunit of β-1, 3-glucan synthase was higher only in intermediate group [3.4(2-8.5), P<0.01]. *HOG*1 MAP kinase showed higher inducible expression in intermediate isolates, while those of *HSP*90-like protein and *Cna*B were comparable in resistant and intermediate groups. In one isolate pan-echinocandin resistance mediated by S639F mutation coupled with high basal chitin content was noted. This study reports novel mutation F635Y/L in *Fks1* in *C. auris*, contributing to echinocandin resistance and suggests the possible adjunctive roles of chitin synthase, *Fks1*, and cell wall-remodeling pathway gene upregulation in caspofungin resistance.

## Introduction

In the last decade, a paradigm shift in the epidemiology of invasive fungal infections has been noticed with frequent isolation of non-*albicans Candida* species from invasive candidiasis (1). Multi-drug resistant *C. auris* has become a significant worry due to a rise in its incidence, outbreak potential, and association with high mortality among hospitalized critically ill patients (2, 3). Since its first report in 2009 from Japan, *C. auris* has been reported in 39 countries with multiple outbreaks ((4–6)).

*C. auris* isolates overwhelmingly exhibit resistance to fluconazole with variable resistance to other triazoles, and the resistance has been found to be clade-specific (7, 8). Considerable resistance to amphotericin B also has been noted in Clade I (South-Asian clade) isolates, which leaves clinicians with no alternative but to use echinocandins as the front-line agents for invasive *C. auris* infections (7, 9). Echinocandins inhibit the activity of 1, 3-*β*-D-glucan synthase, an enzyme that catalyzes the synthesis of primary fungal cell wall polymer, 1, 3-*β*-D-glucan (10). Although many investigational compounds have demonstrated efficacy against *C. auris*, none of those has been approved yet for treatment of *C. auris* (11, 12). ESCMID-ECMM and IDSA clinical practice guidelines recommend echinocandins as first-line treatment options for the management of invasive *Candida* infections (13, 14). However, echinocandin resistance in *C. auris* is also evolving causing further concern while managing *C. auris* infections (7, 15). A study from India reported a high (37%) prevalence of caspofungin resistance in *C. auris* (16), although such high prevalence could be an overestimate confounded by paradoxical growth or ‘eagle effect’ seen mainly with caspofungin. Therefore, it is suggested that elevated caspofungin MIC without an accompanying cross-resistance to other echinocandins should not be treated as a bonafide resistance unless supported by molecular data (15). The echinocandin resistance in *Candida* species is associated with mutations in the *Fks1* gene, which encodes 1, 3-*β*-D-glucan synthase (17). Further, resistance-related mutations are mostly confined to an eight residue motif, the hotspot region 1 of *Fks1* (18). Activation of stress response pathways has been found to play a substantive role in baseline antifungal tolerance and resistance in species of *Candida* and *Aspergillus fumigatus*. The major regulators of stress response pathways include *PKC*, *HOG*1 *MAP* kinase, and Ca^2+^/calcineurin signalling leading to upregulation of chitin synthase genes at the downstream end and *HSP*90 is an essential molecular chaperone that stabilizes these important regulators (19–21).

In the present study, we screened echinocandin resistance in a collection of *C. auris* clinical isolates originating from different centers in India, including six sequential isolates from a single patient. All the isolates with elevated caspofungin MICs were genotyped for mutations in *Fks1*. We evaluated the differential expression of chitin synthase, *Fks*1 and cell wall integrity pathway-related genes in these isolates

## Results

### Antifungal susceptibility testing

Echinocandins susceptibility profile of 199 *C. auris* tested is summarized in Figure 1. The resistant cases were reported from five different centers located in four different provinces of India (Figure 2). MIC90 of all the three echinocandins was 0.5 mg/L. Anidulafungin exhibited better efficacy with geometric mean (GM) MIC 0.18 mg/L compared to caspofungin (GM, 0.38 mg/L) and micafungin (GM, 0.22 mg/L). Seventeen isolates (8.5%) of *C. auris* exhibited elevated caspofungin MIC (≥1 mg/L), with 11 of them also displaying higher MICs (≥1 mg/L) to anidulafungin and micafungin (Table 1). Out of 30 centers included in the study, echinocandin-resistant *C. auris* isolates were reported from five centers. Echinocandin resistance was noted in 6.2% (12/194) of the patients with *C. auris*.

**Figure 1:**
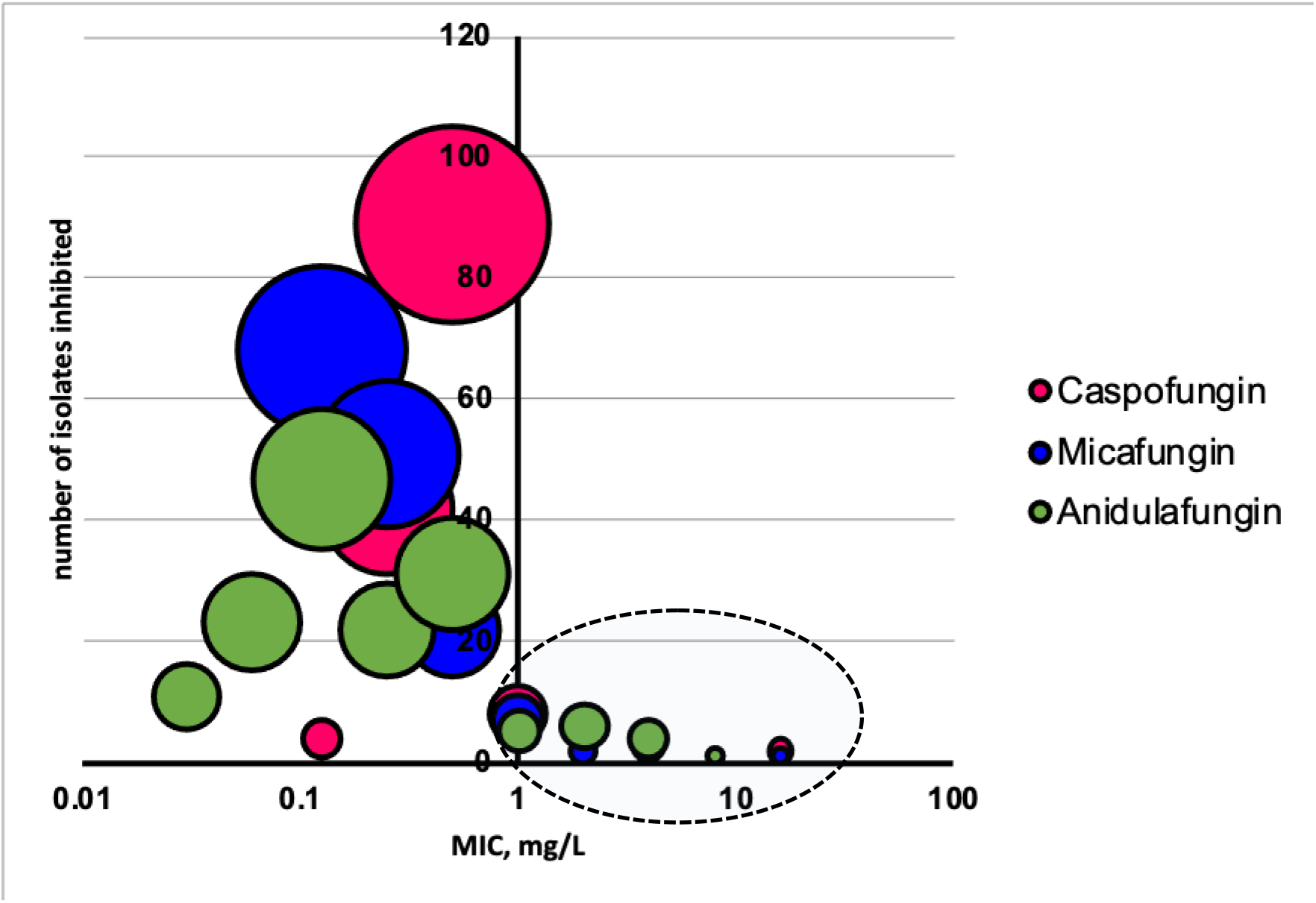
Bubble chart depicting echinocandin MIC distribution profile of 199 *C. auris* isolates determined by broth microdilution method of CLSI M27. The MICs are shown in logarithmic scale and the size of the bubble corresponds with number of isolates inhibited at each MIC. The encircled bubbles indicate isolates with elevated MICs (>1 mg/L)

**Figure 2:**
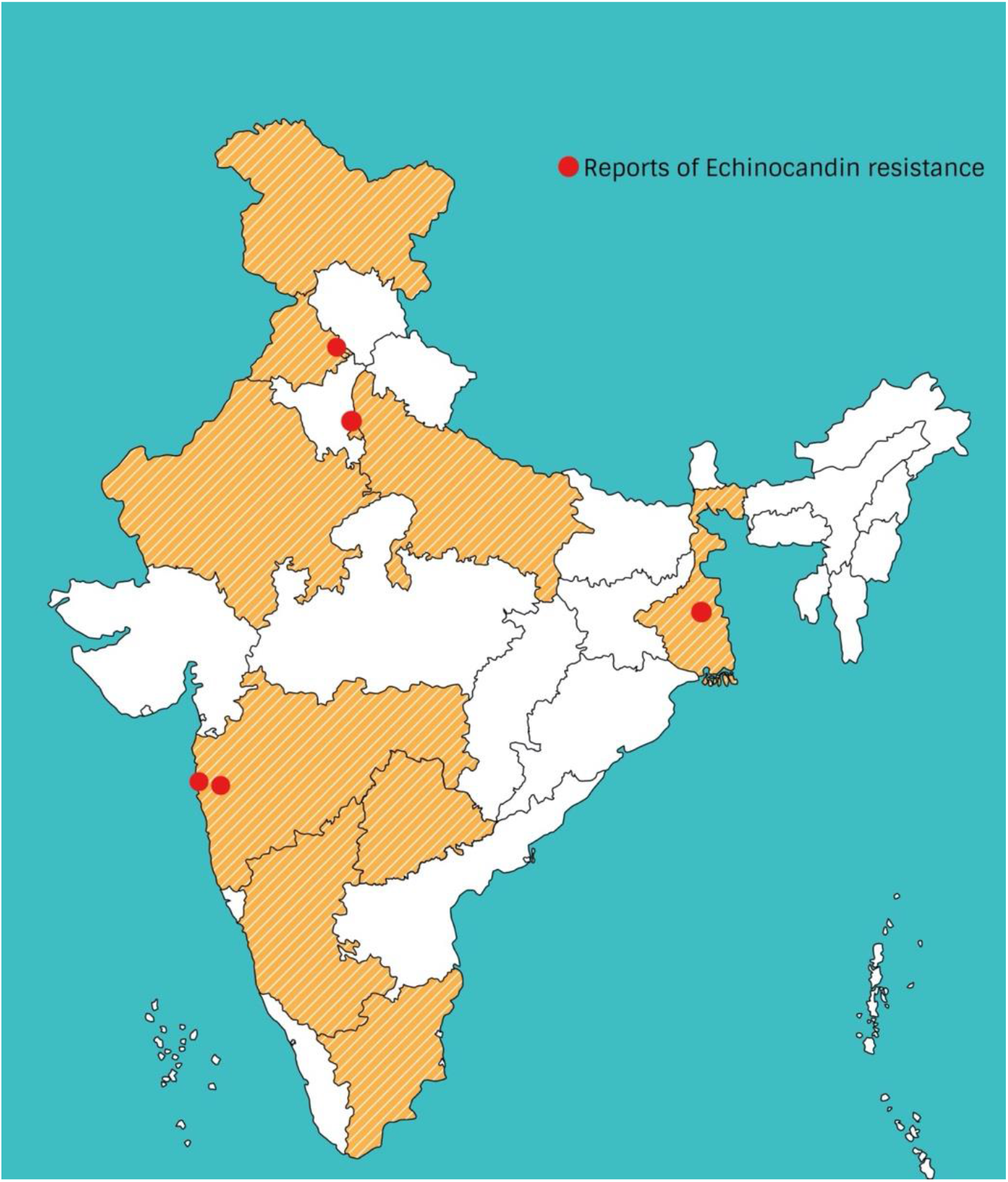
Geographical map of India representing the origin of the isolates used in the study. The dots represent the centres from where echinocandin-resistant isolates of *C. auris* were isolated

**Table 1:** Echinocandin susceptibility profile and *Fks1* genotype of *C. auris* isolates with elevated echinocandin MICs. RM, resistant with non-wildtype *Fks1*; RWT, resistant with wild-type *Fks1*; IWT, isolates with intermediate echinocandin MICs and wild-type *Fks1*; SWT, susceptible isolates with wild-type *Fks1*; BALF, bronchoalveolar lavage fluid; TTS, tracheal tube secretion

|  |  |  |  | MIC (mg/L) |  |  |
| --- | --- | --- | --- | --- | --- | --- |
| Isolate ID | Sample | Phenotype | *FksI* genotype | CAS | AFG | MFG |
| NCCPF470197 | Blood | Resistant | T1904A(F635Y) | 16 | 1 | 1 |
| NCCPF470198 | Urine | Resistant | C1916T(S639F) | 4 | 4 | 1 |
| NCCPF470199 | Urine | Resistant | C1916T(S639F) | 4 | 4 | 1 |
| NCCPF470203 | Blood | Resistant | C1916T(S639F) | 2 | 8 | 16 |
| NCCPF470209 | Blood | Resistant | T1904A(F635Y) | 4 | 2 | 1 |
| NCCPF470237 | Urine | Resistant | T1903C(F635L) | 1 | 2 | 1 |
| NCCPF470238 | Urine | Resistant | T1903C(F635L) | 1 | 2 | 1 |
| NCCPF470240 | Blood | Resistant | T1903C(F635L) | 1 | 2 | 0.5 |
| NCCPF470241 | TTS | Resistant | T1903C(F635L) | 1 | 2 | 1 |
| NCCPF470200 | Urine | Resistant | WT | 16 | 2 | 2 |
| NCCPF470201 | Blood | Resistant | WT | 2 | 1 | 2 |
| NCCPF470239 | BALF | Intermediate | WT | 0.5 | 1 | 0.25 |
| NCCPF470242 | TTS | Intermediate | WT | 0.5 | 1 | 0.25 |
| NCCPF470202 | Blood | Intermediate | WT | 1 | 0.5 | 0.5 |
| NCCPF470204 | Blood | Intermediate | WT | 1 | 0.5 | 0.5 |
| NCCPF470206 | Blood | Intermediate | WT | 1 | 1 | 0.25 |
| NCCPF470205 | Blood | Intermediate | WT | 1 | 0.5 | 0.5 |
| NCCPF470196 | Blood | Susceptible | WT | 0.12 | 0.06 | 0.12 |
| NCCPF470208 | Blood | Susceptible | WT | 0.12 | 0.25 | 0.25 |
| NCCPF470210 | Blood | Susceptible | WT | 0.12 | 0.25 | 0.25 |
| CBS12372 | Blood | Susceptible | WT | 0.12 | 0.12 | 0.25 |

### *Fks1* sequence analysis

Seventeen isolates with variable resistance to caspofungin were subjected to *Fks1* sequence analysis; nine isolates exhibiting MIC ≥2 mg/L for any of the three echinocandins carried an acquired mutation in the HS1 region (Table 1). Among those nine isolates, three harboured S639F mutation, which has been previously implicated in echinocandin resistance in *C. auris*. However, a novel mutation, F635Y, was found in two isolates with caspofungin MICs of 4 and 16 mg/L. Of six sequential isolates from a single patient, four had F635L substitution resulting in elevated MICs to anidulafungin. Two other isolates with caspofungin MIC of 2 mg/L and 16 mg/L showed no alteration in HS1 region of *Fks1*. Six isolates exhibiting MIC of 1 mg/L for caspofungin or anidulafungin also carried a wild-type *Fks1* HS1 genotype. The *Fks1* nucleotide sequence of nine non-wild type isolates were submitted to the GenBank with the accession numbers ranging from MT199096-MT199102, MT199103 and MT199104.

### Expression analysis of chitin, *Fks1* and cell wall stress response genes

On induction with respective sub-MIC caspofungin concentration, both resistant and intermediate isolates demonstrated significantly higher (*P*<0.05) induction of *Chs*1 [resistant, 2.2 (1.3-5.8); intermediate 6 (2.5-11.2)] compared to susceptible comparator group [1(0.8-2.0)] (Figure 3A). Also, one among three isolates in the resistant group with S639F showed higher induction of the *Chs*1 gene (8.3±1.8) compared to the other two isolates (*P*=0.0001) (Figure 3B). Similarly, both resistant and intermediate groups demonstrated significantly higher (P<0.05) upregulation of *Chs*2 [resistant, 3(2.2-4); intermediate, 6.8(2.3-13.5)] compared to susceptible isolates [1.2(0.9-1.4)] (Figure 3C).

**Figure 3:**
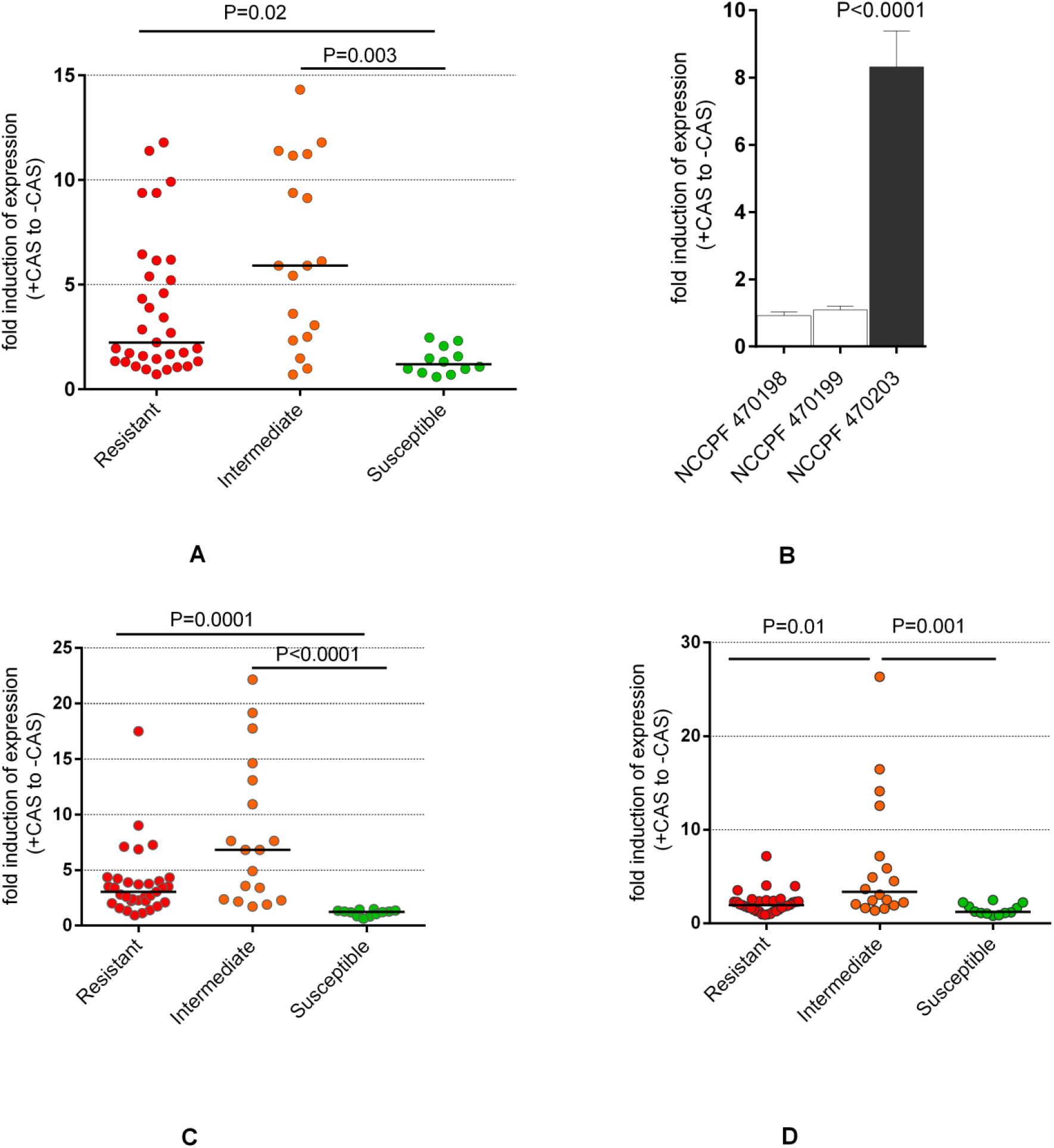
Fold-induction in expression of putative chitin synthase gene homologues, (A) *Chs*1 and (C) *Chs*2, and (D) *Fks1*. The median fold changes represent values from experiments put up at three different time points. The data was analysed using Kruskal-Wallis test with Dunn’s post-hoc test for multiple comparisons. Figure 3B represents fold-induction of *Chs*1 in three resistant isolates with S639F *Fks1* mutation. The data was analysed using one way ANOVA with Bonferroni’s test for multiple comparisons

A differential transcriptional upregulation of *Fks1* gene was also observed across the groups (P<0.0001) (Figure 3D). However, only intermediate group isolates demonstrated higher induction [3.3 (2-8.5)] of *Fks1* compared to both susceptible [1.2 (1.0-2.0), P<0.01] and resistant group [2(1.4-2.4), P<0.05].

Among the cell wall stress response genes, expression of the gene encoding *HSP*90-like protein was higher in resistant [2(1-3), P<0.001] and intermediate isolates [3.5(1.7-12), P<0.0001] compared to susceptible isolates [0.7(0.4-1.0)], but comparable expressions levels were noted in resistant and intermediate groups (P>0.05) (Figure 4A). While *HOG*1 MAP kinase demonstrated higher induction in isolates with intermediate susceptibility [3.72(1.96-8.2)] compared to resistant [2(1.2-2.8), P<0.01] as well as susceptible isolates [1.2(1-1.4), P<0.001] (Figure 4B). The expression of calcineurin B subunit (*Cna*B) was low in all the three groups [R, 0.9(0.7-1.0); I, 0.8(0.7-0.85); S, 1.1 (0.9-1.2)] but feebly higher in susceptible isolates (P<0.01) (Figure 4C).

**Figure 4:**
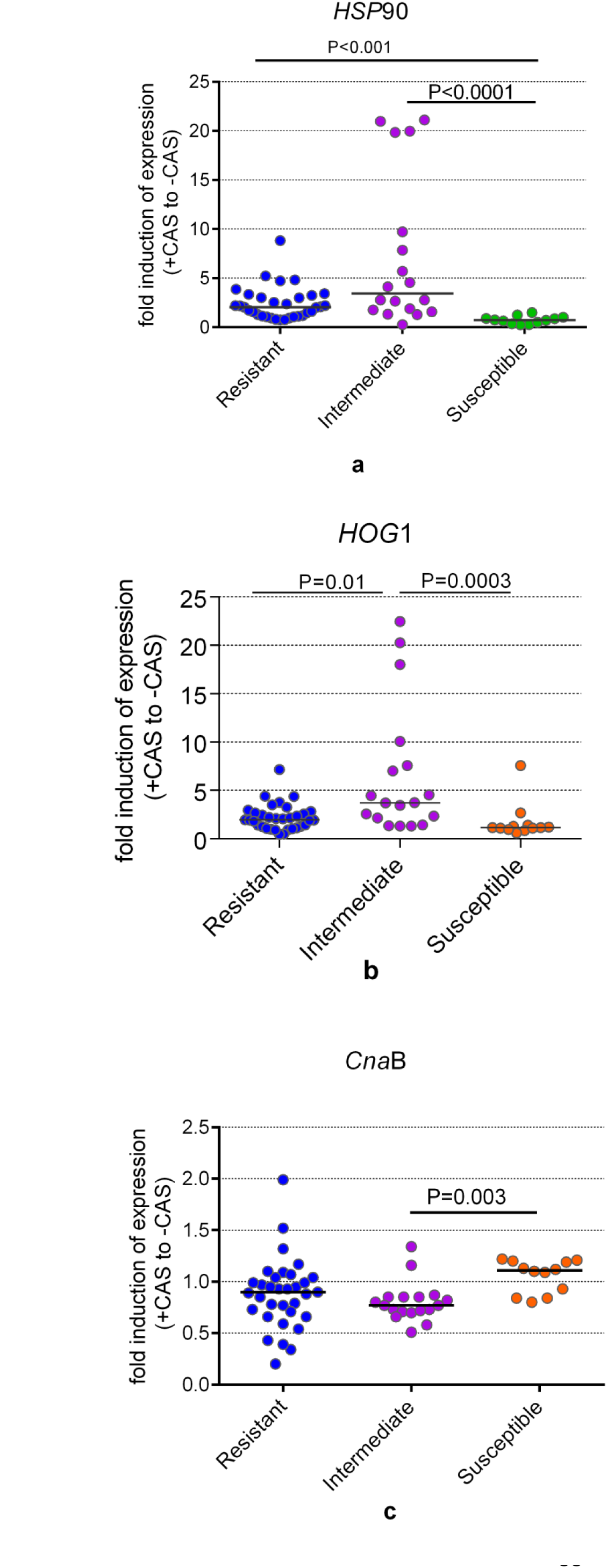
Fold-induction in expression of cell wall stress response pathway genes, including HSP90-like protein (a), HOG1 MAP kinase (b), and calcineurin subunit B (c). The median fold changes represent values from experiments put up at three different time points. The data was analysed using one-way Kruskal-Wallis test with Dunn’s post-hoc test for multiple comparisons

### Chitin content quantification

The baseline and inducible chitin content were evaluated by the flowcytometry-based quantification method. The baseline chitin contents [median staining index (interquartile range)] did not vary across the three groups: Resistant, 9.6 (2.8-20.2); Intermediate, 10.5 (2.6-16); susceptible 17 (14-24) (P>0.05) (Figure 5). However, exposure to caspofungin resulted in a differential increase in chitin levels in resistant and intermediate isolates compared to their untreated controls (P<0.0001). On the other hand, the chitin levels were not induced significantly in susceptible isolates (P>0.05). Bland-Altman analysis demonstrated close agreement between *Chs*1 fold-changes and the staining index (Figure 6). The bias between *Chs*1 drug-inducible transcriptional levels and inducible chitin content was 0.21 [95% C.I. (−0.22 to 0.65)].

**Figure 5:**
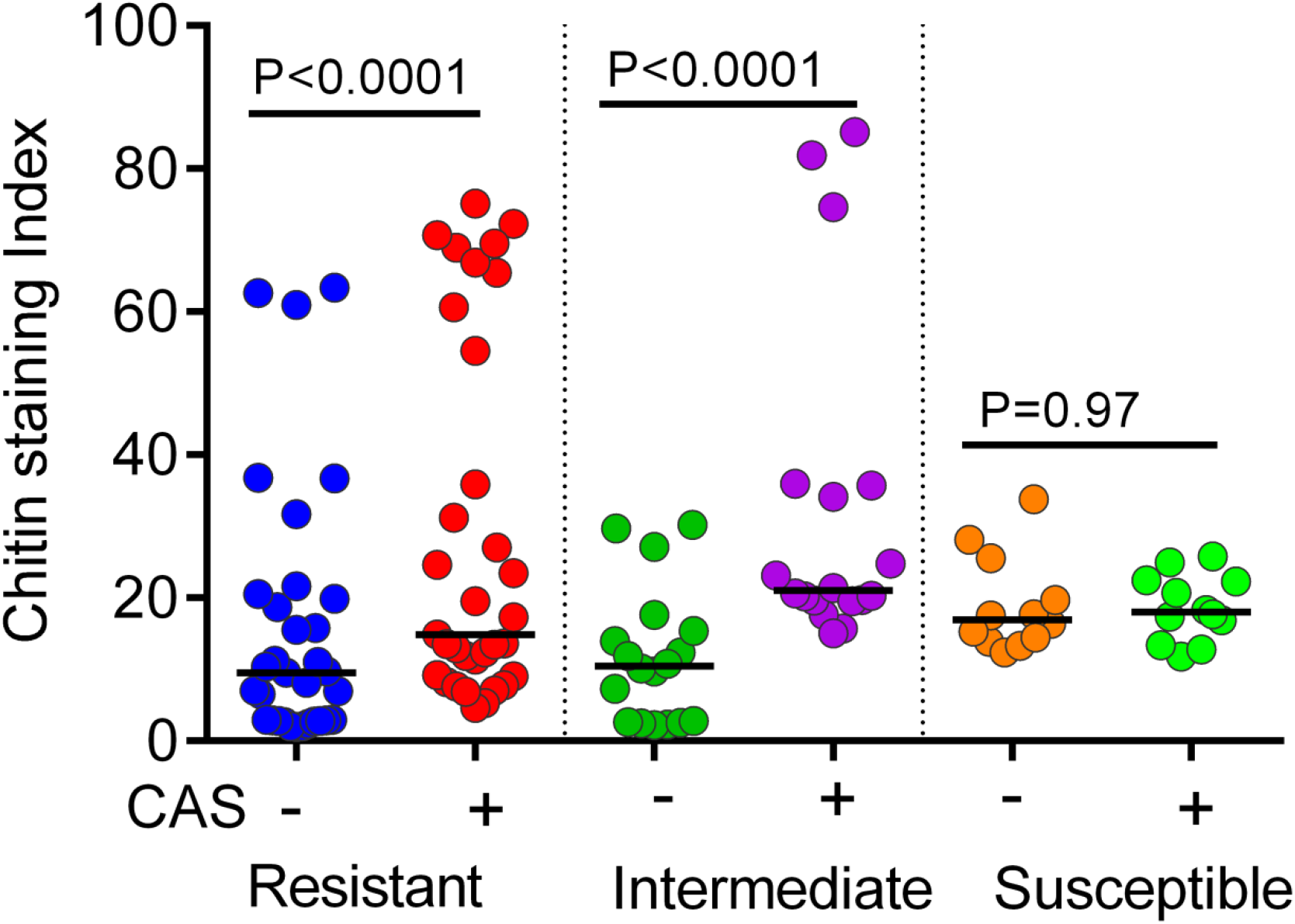
Baseline and caspofungin-induced differential chitin content represented as staining index. Three biological replicates were used for each isolate to determine the chitin levels and each pair of data were analyzed by Wilcoxon rank test

**Figure 6:**
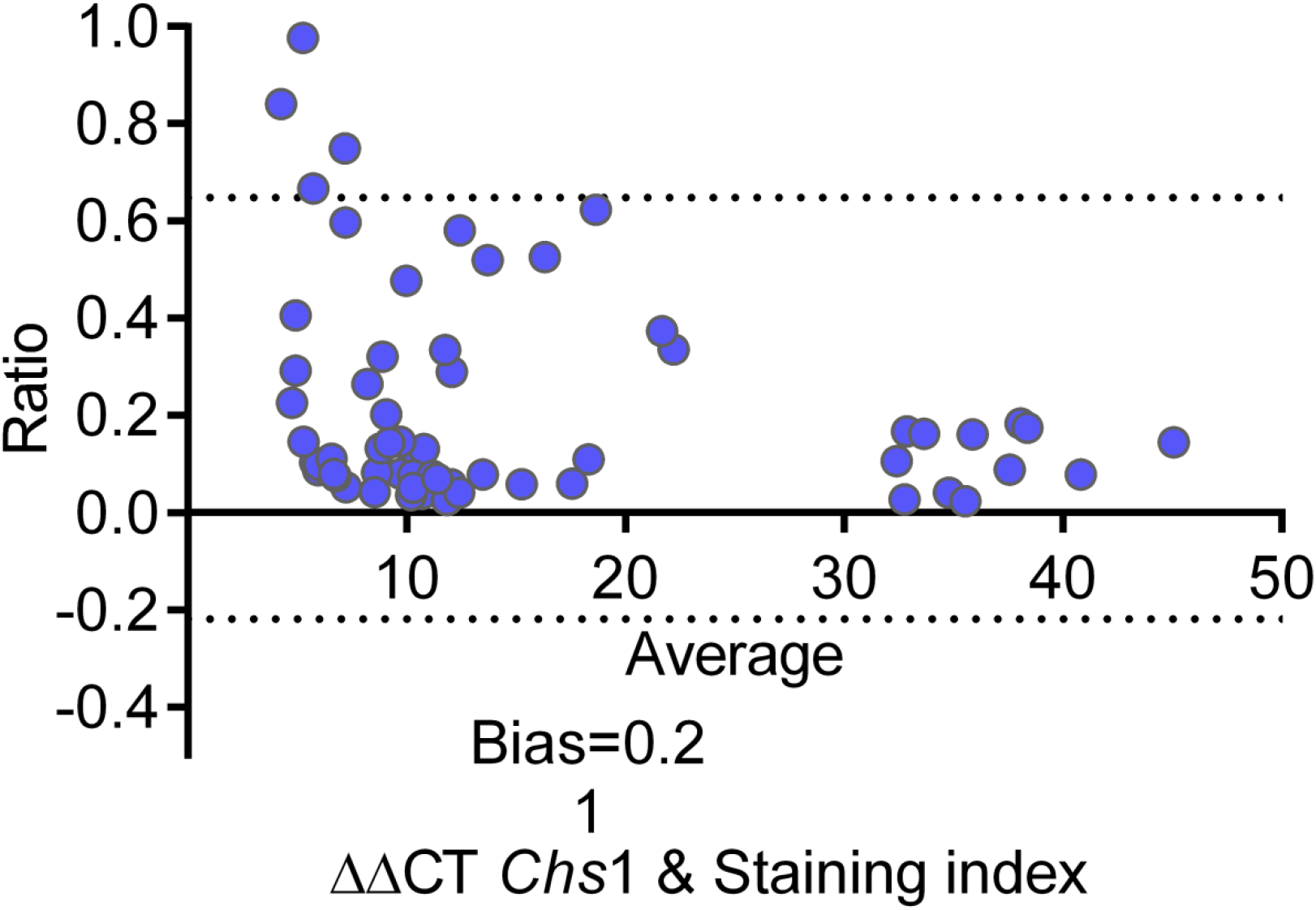
Bland-Altman analysis demonstrating agreement between transcriptional fold changes in *Chs*1 and the staining index values. The dotted lines denote 95% limits of agreement between the two measurement methods. The bias is calculated as the difference between values determined by two assays and proximity of this value to zero indicates good agreement between the two assays.

### Growth Kinetics

Growth curve analysis demonstrated synchronous growth pattern of resistant, intermediate, and susceptible isolates with no differential growth rates (Figure 7). The *Fks*1 genotype or echinocandin susceptibility of the isolate did not impact the phased growth curves of the isolates (*P*=0.44).

**Figure 7:**
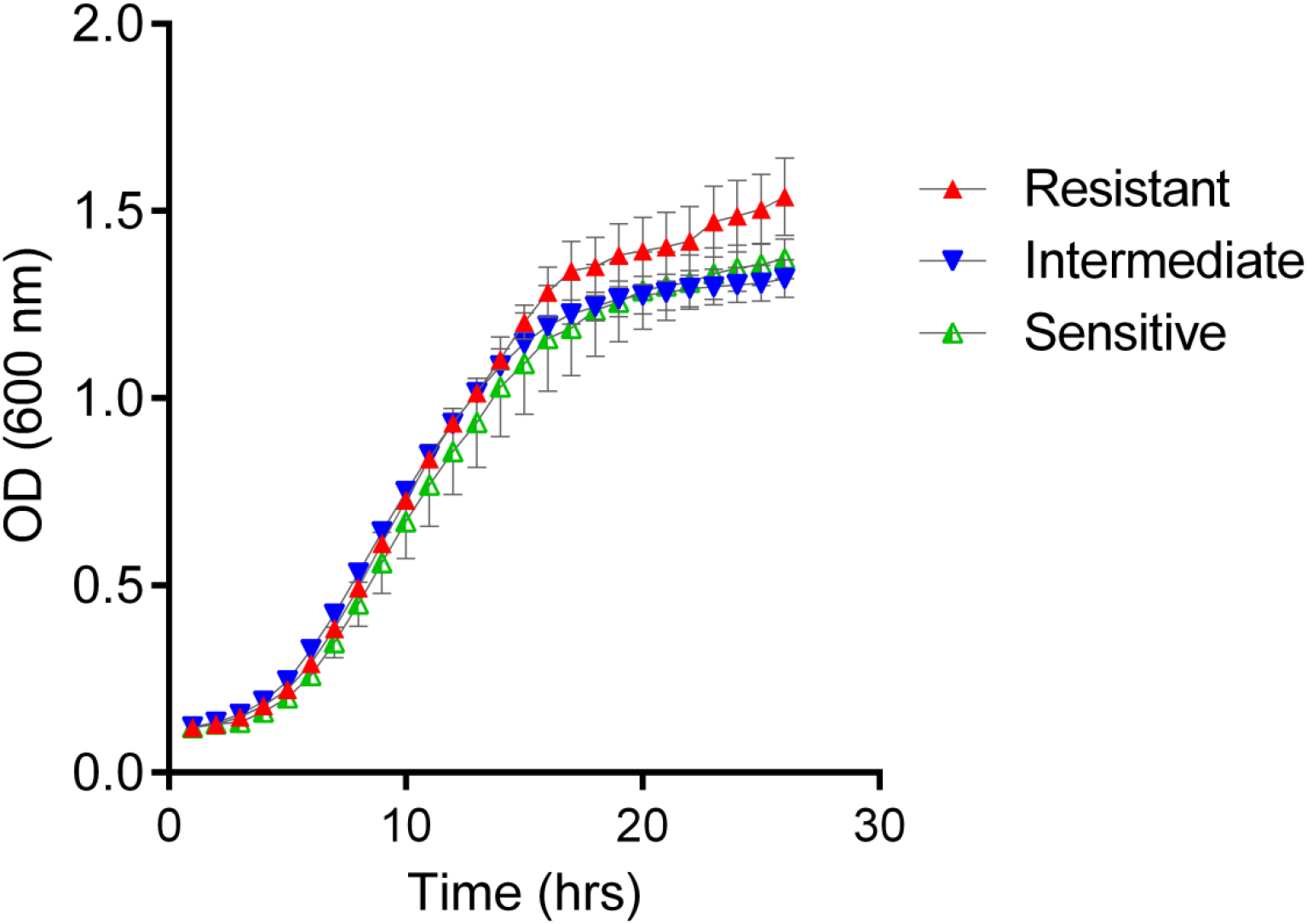
Growth curve of *C. auris* isolates represented as mean OD values ±SE of R (*n*, 6), I (*n*, 4), and S (*n*, 4) isolates at each 1 hr intervals throughout a 24 hr growth period.

### Clinical details of the patient with six serial isolates

Six sequential *C. auris* were isolated from urine (*n*, 2) blood (*n*, 1), bronchoalveolar lavage fluid (BALF; *n*, 1), and tracheal tube secretions (*n*, 2) from a 37-year-old, male patient with liver transplantation. Transplantation was performed due to alcoholic liver disease and cirrhosis (Figure 8). Post-transplant underwent splenectomy, portocaval shunting, laparotomy for bleeding, and biliary stenting for cholangitis. Subsequently the patient developed severe manifestations of thrombotic thrombocytopenic purpura and underwent central venous and indwelling urinary catheterization, and placement of the tracheal tube. Given multiple surgical interventions & immunosuppressive conditions, the patient was put on Liposomal amphotericin B prophylaxis at 3 mg/kg daily, as per standard protocol of the hospital. During antifungal prophylaxis, an echinocandin-resistant *C. auris* with an *Fks1* mutation (#1 isolate) was isolated from his urine. Subsequently, he received a combination therapy of posaconazole intravenously 400 mg 1^st^ day, followed by 200 mg daily and micafungin 150 mg daily for suspected additional lung shadows. Despite micafungin therapy the candiduria due to *Fks1* mutant echinocandin-resistant *C. auris* (#2 isolate) persisted. Posaconazole was withdrawn on day 4 of the combination therapy due to deranged liver function, but micafungin was continued. On the same day of posaconazole withdrawal, broncho-alveolar lavage (BAL) sample yielded a wild-type isolate of *C. auris* (#3 isolate). Next day, echinocandin-resistant *C. auris* with *Fks1* mutation was isolated from blood and tracheal aspirate (#4 and #5 isolate). As there was no alternative antifungal treatment available, micafungin was continued. Subsequently after a week of blood isolation, microbiological clearance of candidemia was achieved. However, a new WT *C. auris* strain (#6 isolate) was isolated from tracheal tube secretions at the same time, and no clinical improvement of the patient was noticed. Finally, the patient developed recurrent bleeding and thrombosis due to refractory thrombotic thrombocytopenic purpura and left the hospital against medical advice and died soon after. The antifungal susceptibility profile of these isolates is provided in table 2. AFLP analysis of the six isolates demonstrated that three isolates recovered from lower respiratory tract specimens (one BAL fluid and two from tracheal tube secretion) formed one cluster with fingerprint similarity 98.5% (Figure 9). Two isolates recovered from urine samples formed a second cluster (98% fingerprint similarity), while the blood isolate was different from the above two clusters.

**Figure 8:**
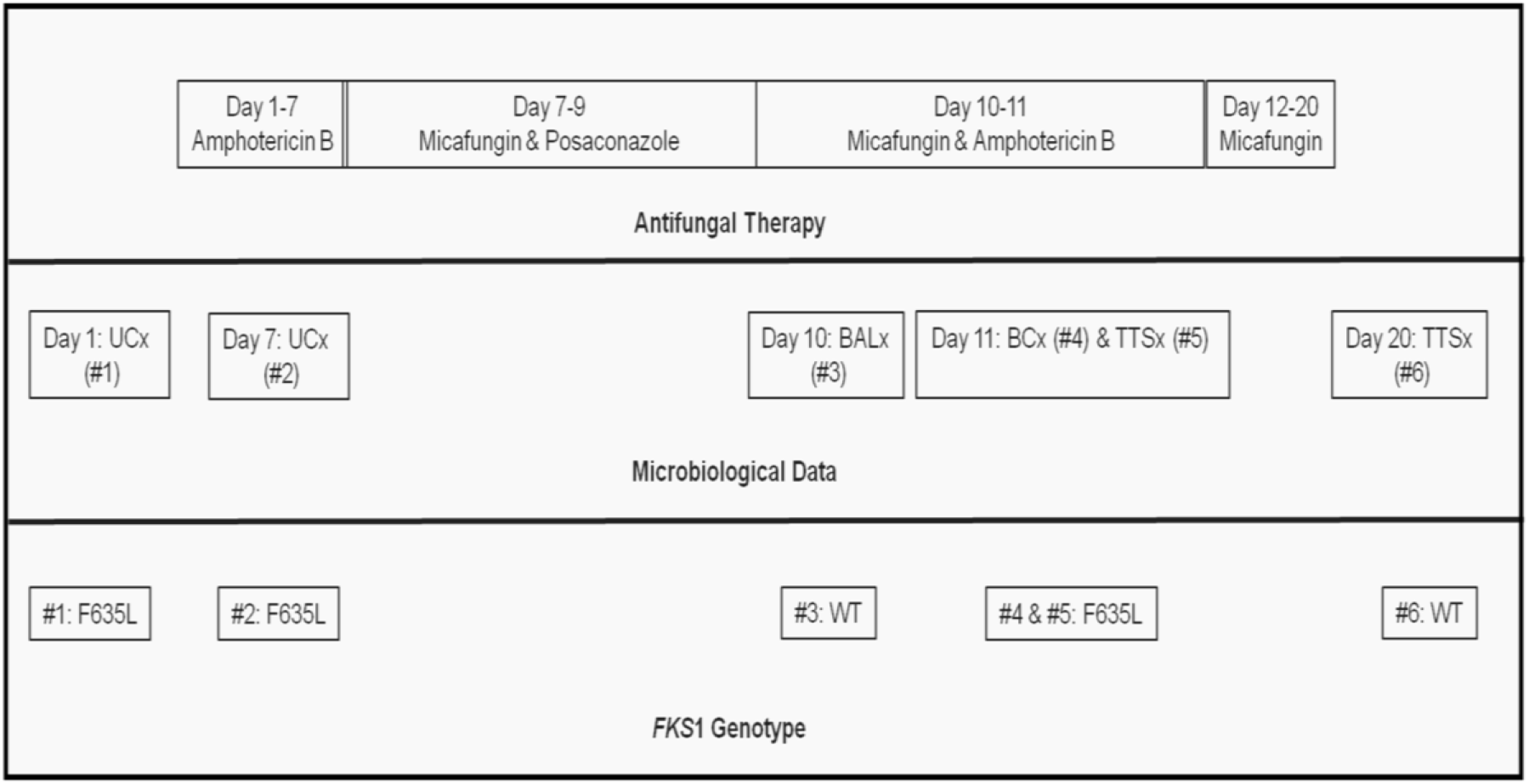
Timeline of antifungal therapy and microbiologic data of a liver transplant patient with six sequential *C. auris* isolates and their *Fks1* genotype

**Figure 9:**
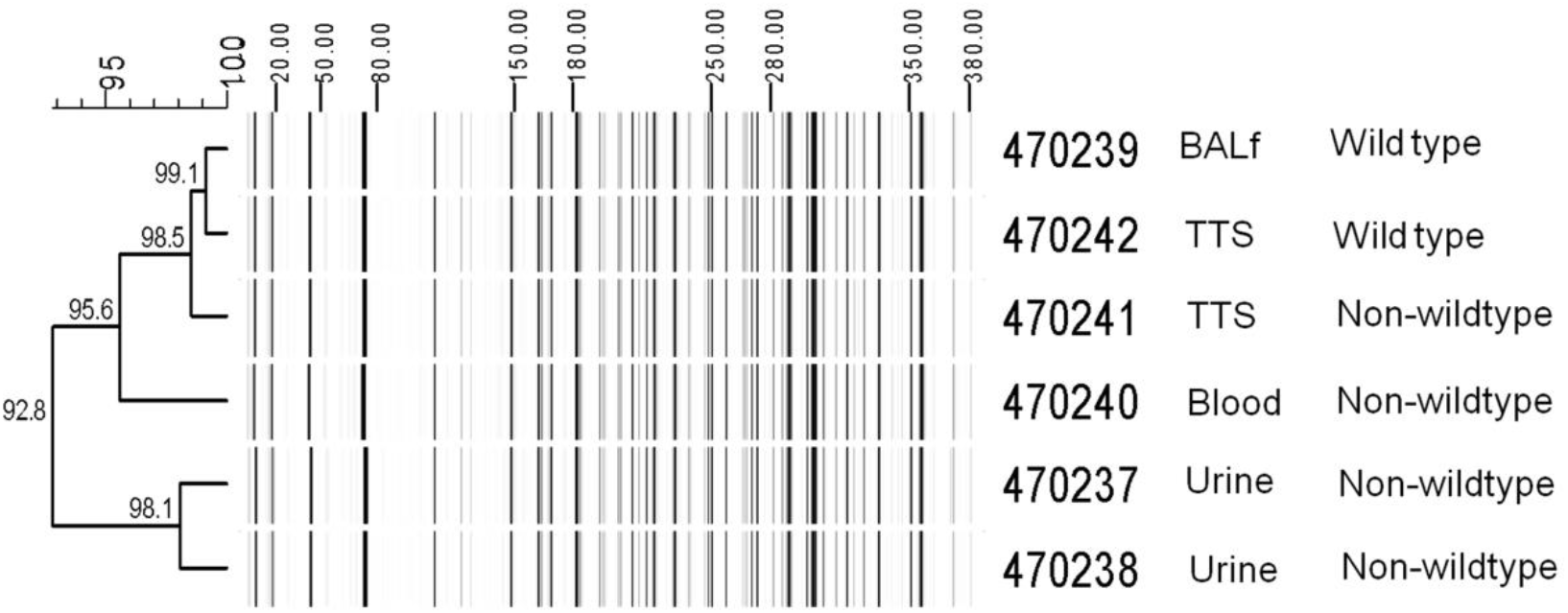
Amplified fragment length polymorphism analysis of 6 sequential isolates of *C. auris*. BALf, bronchoalveolar lavage fluid; TTS, tracheal tube secretion

**Table 2:** Antifungal susceptibility profile and *Fks1* genotype of six sequential *C. auris* isolates with their sources and dates of isolation

| S. No. | NCCPF ID | Date of Isolation | Sample | AMB | FLC | VRC | ITC | PSC | CAS | AFG | MFG | *Fks1* genotype |
| --- | --- | --- | --- | --- | --- | --- | --- | --- | --- | --- | --- | --- |
| #1 | NCCPF470237 | 18/11/2019 | Urine | 1 | 16 | 0.125 | 0.03 | 0.03 | 1 | 2 | 1 | T1903C(F635L) |
| #2 | NCCPF470238 | 24/11/2019 | Urine | 1 | 16 | 0.125 | 0.03 | 0.03 | 1 | 2 | 1 | T1903C(F635L) |
| #3 | NCCPF470239 | 27/11/2019 | BALF | 1 | >64 | 1 | 0.25 | 0.25 | 0.5 | 1 | 0.25 | Wild Type |
| #4 | NCCPF470240 | 28/11/2019 | Blood | 1 | 8 | 0.125 | 0.06 | 0.03 | 1 | 2 | 0.5 | T1903C(F635L) |
| #5 | NCCPF470241 | 28/11/2019 | TTS | 1 | 16 | 0.125 | 0.06 | 0.03 | 1 | 2 | 1 | T1903C(F635L) |
| #6 | NCCPF470242 | 7/12/2019 | TTS | 1 | >64 | 1 | 0.25 | 0.25 | 0.5 | 1 | 0.25 | Wild Type |

## Discussion

Echinocandins are frontline drugs in the management of azole-resistant invasive *C. auris* infections. Therefore, the emergence of echinocandin resistance in *C. auris* is a matter of great concern. In the present study, 8.5% (17/199) of *C. auris* isolates had elevated echinocandin MICs, though true resistance as per CDC recommended cut-off values was 5.5% (11/199). It is pertinent to mention here that CDC has derived these breakpoints from extrapolation of susceptibility and PK/PD data of related species, and not from that of *C. auris*. Therefore, determination or the revision of existing MIC cut–off values for echinocandin susceptibility of *C. auris* would be important, as four of our isolates had anidulafungin MIC one to two-fold dilution lower than the proposed cut-off (≥4 mg/L), but those isolates carried a mutation, F635L in *Fks1*. Similarly, two isolates (NCCPF470200, NCCPF470201) exhibited higher MICs for caspofungin with a wild type *fks*1 genotype. Considering the presence of a genetic marker of echinocandin resistance, 4.5% (9/199) of our isolates would qualify as resistant to at least one echinocandin. Other series, including our previous report on echinocandin resistance in *C. auris* also reported similar (2-9.5%) resistance rates (7, 8, 15, 22).

*C. auris* is known for horizontal transmission in clinical settings owing to its persistence in the healthcare environment and acquisition from other patients, environment and rarely from healthcare workers (3, 23). In the absence of a prophylactic or empiric echinocandin therapy, development of echinocandin–resistant *C. auris* infection in multiple cases would suggest a single source and clonal origin of the resistant isolates. However, none of the patients with resistant *C. auris* in the present series was administered either prophylactic or empiric echinocandin therapy before the isolation of *C. auris*. In the patient with six serial heterogeneous populations of *C. auris*, the first caspofungin-resistant isolate was obtained when the patient was on amphotericin B therapy (Figure 8). Also, all the resistant cases were reported from five different centers of India with bleak chances of a single source acquisition (Figure 2).

Acquired resistance in *Candida* species to echinocandins was first reported in 2005 with mutations in 1,3-*β*-D-glucan synthase subunit, *Fks1.* The mutations impart resistance, or reduced susceptibility to echinocandins (17). Point mutations leading to amino acid substitutions in the hot spot regions of *Fks1* subunit conferring echinocandin resistance had been reported in other *Candida* species ((24–26)). *S. cerevisiae* possesses two paralogs of the *FKS* gene, *Fks1*, and *Fks*2, which are differentially regulated. In *Fks1*Δ mutants of *S. cerevisiae*, *Fks*2 takes over the role of *Fks1*, and similar echinocandin resistance-associated point mutations arise in *Fks*2. Likewise, *C. glabrata* has been reported to carry three syntenic orthologs, *Fks1*, *Fks*2, and *Fks*3. However, echinocandin resistance-related mutations in *C. glabrata* have been found more frequently in *Fks*2 than *Fks1*. In *C. auris* there is a single orthologue of *Fks1*, but any paralog of this gene has not been reported (7, 15, 27). Both experimental and clinical data have demonstrated therapeutic failure and increased all-cause mortality associated with a mutation in *the Fks1* gene in echinocandin resistance (15, 24). In the present study, of the 11 isolates with bonafide resistance, nine isolates harbored a mutation in the *Fks1* gene, among which two isolates had F635Y genetic alteration. Despite carrying the same mutation, these two isolates demonstrated a varying degree of resistance, with one isolate exhibiting high-level resistance to caspofungin (MIC, 16 mg/mL) compared to the other isolate (MIC, 4 mg/mL). Previously, only S639F was known to confer echinocandin resistance in *C. auris* (7, 15). It is generally claimed that in the absence of this mutation, high MIC to echinocandin may not be actual resistance. However, other molecular mechanisms of resistance may exist. In the present study, a novel F635Y/L mutation conferred resistance in 6 isolates. This non-synonymous substitution, F635Y/L, occurs at the residue marking the beginning of the hotspot 1 region of the *Fks1* subunit in *C. auris* and its equivalence in *C. albicans* rests on F641Y(15, 28). The mutation, F635L, was demonstrated in four isolates in the present study. In *C. glabrata*, the mutation at this position leads to the change of amino acid, which imparts varying degrees of resistance (29). In our study, the substitution of phenylalanine at position 635 with leucine in those four isolates conferred different susceptibility pattern from F635Y. The phenylalanine substitution led to high anidulafungin MIC (2 mg/L), but modest increase in the MIC of caspofungin and micafungin (1 mg/L each).

Echinocandins blunt the ability of 1, 3-*β*-D-glucan synthase to synthesize optimum amounts of cell wall polymer, 1, 3-*β*-D-glucan, which results in cell wall stress. In response to this cellular stress, cells activate many adaptive salvage mechanisms (18). Chitin and 1, 3-*β*-D-glucan are the two crucial cell wall polymers described as the ‘yin and yang’ of the fungal cell wall (30). Compensatory induction of chitin synthesis is an essential adaptive stratagem to ameliorate the cell wall stress imposed by echinocandins (18). In *C. albicans*, *PKC*, *HOG1 MAP* kinase, and Ca^2+^/calcineurin signalling circuitry regulate this cell wall salvage pathway (19). *HSP9*0 is a hub of cell signalling pathways that stabilizes crucial regulators of the cellular stress response. *HSP*90 promotes basal antifungal drug tolerance and resistance to both azoles and echinocandins. Pharmacological or genetic inhibition of *HSP*90 has been demonstrated to abrogate resistance in *C. albicans* and *A. fumigatus* (21). In the present study, the intermediate group exhibited higher induction of *HSP*90-like protein and *HOG*1 MAP kinase genes, whereas resistant isolates demonstrated higher upregulation of *HSP*90 only. Surprisingly, the inducible expression of calcineurin regulatory subunit, *Cna*B, was low in all the isolates, though susceptible isolates showed slightly higher expression.

The absence of an *Fks1* mutation in two resistant and six intermediate isolates with no known functionally annotated paralog of *Fks1* in *C. auris* prompted us to evaluate the differential gene expression of *Fks1* and two putative chitin synthase genes, *Chs*1 and *Chs*2.

Exposure of *C. albicans* to low levels of echinocandin has been shown to stimulate chitin synthase gene expression resulting in elevated chitin synthesis and increased tolerance to echinocandins (31). We also observed that inducible expression of chitin synthase genes, *Chs*1 and *Chs*2, were relatively higher in resistant and intermediate isolates than in susceptible isolates. However, the inducible upregulation of *fks1* was higher in only the intermediate group compared to susceptible isolates. We believe that intermediate isolates with elevated caspofungin MICs but short of one two-fold dilution from the suggested cut-off value for resistance (≥2 mg/L), are intermediary populations resulting from activation of stress response pathways. The isolates of this kind failed to induce therapeutic failure in murine pharmacodynamic models but were presumed to be a transitional phase in the evolution of bonafide resistance (18).

Three resistant isolates with S639F mutation could easily be delineated based on their differential echinocandin susceptibility pattern. While two of those isolates had identical susceptibility patterns, one isolate NCCPF 470203, exhibited high-level MICs to anidulafungin and micafungin. The inducible levels of the *Chs*1 gene and chitin content in this isolate was higher compared to rest of the isolates in the R category (Figure 3B). Pertinently, mutation in *Fks1* coupled with high chitin contents was found associated with echinocandin resistance in *C. albicans* (32).

Exposure with corresponding sub-inhibitory caspofungin concentrations led to a significantly higher upregulation of *Chs*1 and *Chs*2 genes in resistant and intermediate isolates. Also, the inducible expression of *Fks1* in intermediate isolates was significantly higher than either of the other two groups, which indicated that decreased susceptibility in those two groups of isolates could be due to the transient upregulation of chitin synthase and/or *Fks1* genes. However, since both resistant and intermediate isolates had comparable upregulation in Chitin synthase and *Fks1*, the transient alterations in the chitin and 1, 3-*β*-D-glucan synthase genes were unlikely to account for the high-level resistance demonstrated by two resistant isolates.

AFLP analysis of isolates sequentially recovered from a liver transplant patient showed more than 92% similarity in their banding pattern, suggesting that they could be the clonal origin. Given the absence of uniform criteria for the percentage similarity to label it as clonal, the possibility of heterogeneous subpopulations implying multimodal acquisition of *C. auris* by the patient could not be excluded. A recent study also demonstrated the existence of heterogeneous populations of echinocandin-resistant and susceptible *C. auris* isolates in single patients (33).

In *C. albicans* and *C. glabrata*, a trade-off between *Fks*1/*Fks*2 mutations and fitness of the organism has been reported in both clinical and experimental populations of the yeast (34, 35). Notably, increased chitin content in *Fks*1 mutants was associated with attenuated growth translating to hypovirulence in animal models of disseminated infection (34). In the present study, though both resistant and intermediate isolates had higher induction of chitin, none of the *Fks*1 mutation was associated with impaired growth rate and all the three groups of isolates showed similar growth rates under ambient *in vitro* conditions. However, the fitness of these *Fks*1 mutants needs evaluation in animal models.

In conclusion, our study demonstrates that echinocandin resistance is emerging among Indian *C. auris* clinical isolates. We also report novel mutations, F635Y/L, in *Fks1* conferring echinocandin resistance. Further, a high baseline chitin level in conjunction with an *Fks1* mutation may impart a high-level cross-echinocandin resistance. The study also suggests the adjunctive roles played by transcriptional upregulation of chitin synthesis and cell wall stress response genes in promoting echinocandin resistance in *C. auris*.

## Materials and methods

### Isolates and growth conditions

A total of 199 *C. auris* isolates from 194 invasive candidiasis cases from 30 centers of the country during September 2017 through March 2020 were evaluated in this study. The isolates are part of a nation-wide antimicrobial resistance surveillance program done under the auspices of Indian Council of Medical Research, New Delhi (https://www.icmr.nic.in/sites/default/files/guidelines/candida_Auris.pdf) and stored at nodal co-ordinating center, Postgraduate Institute of Medical Education and Research, Chandigarh as per the advisory of Indian Council of Medical Research, New Delhi (https://www.icmr.nic.in/sites/default/files/guidelines/candida_Auris.pdf). The isolates were retrieved, and the identity of each isolate was confirmed by matrix-assisted laser desorption ionization-time of flight (MALDI-TOF) mass spectrometry (Bruker Daltonic GmbH, Bremen, Germany) as per previously described protocol (36).

### Antifungal susceptibility testing

Antifungal drug susceptibility testing was done according to the CLSI broth microdilution method following the M27-protocol (37). *C. parapsilosis* ATCC22019 and *C. krusei* ATCC6258 were used as quality control strains. For azoles (azole susceptibility tested for 6 sequential isolates only) and echinocandins, drug concentrations inhibiting 50% visible growth and for amphotericin B 100% growth inhibition were taken as minimum inhibitory concentration (MIC). The susceptibility data were interpreted as per tentative breakpoints suggested by Centre for Disease Control and Prevention (CDC), Atlanta (38).

### ITS and *Fks1* gene sequencing

Genomic DNA was extracted using phenol-chloroform-isoamyl alcohol according to an earlier described method (39). All isolates having higher MIC (≥1mg /L) to any echinocandin and four echinocandin sensitive isolates were used for further analysis. The identification of those isolates was confirmed by sequencing internal spacer (ITS) region of rDNA. The primer pairs (*Cau*_HS1-F, 5-GCCATCTCGAAGTCTGCTCA-3; *Cau*_HS1-R 5-TGACAATGGCATT CCACACCT-3) were designed to amplify hotspot region-1(HS1) of *Fks1* gene (XM_018312389.1) using the NCBI primer blast tool (https://www.ncbi.nlm.nih.gov/tools/primer-blast/). PCR cycling conditions included initial denaturation at 94 °C for 2 min, with 35 cycles of the program 94 °C for 15 s, annealing at 60 °C for 30 s, and a final extension at 68 °C for 1 min. Gene sequencing was performed using the BigDye terminator ready reaction kit (version 3.1; Applied Biosystems, Foster City, CA, USA) in ABI Genetic Analyzer 3500Dx (Applied Biosystems, Foster City, CA, USA). The raw sequences were analyzed using Seqman software (DNA Star, Laser Gene, ABI) for any nucleotide substitutions. Amino acid sequences were obtained using an online ExPASy tool (http://www.expasy.org/translate) and were aligned in ClustalX2 software.

### Gene expression analyses

To quantify the transcript levels, primers were designed for sequences corresponding to *Fks1*, two putative chitin synthase genes, *Chs*1, *Chs*2; cell wall stress response pathway genes, *HSP*90-like protein, *HOG*1 MAP kinase and *Cna*B subunit. The housekeeping gene, actin, was used as a reference for normalization of the Ct values (Table 3). RNA extraction was done using Trizol (Ambion Life sciences). Briefly, 1×10^6^ cells were seeded into YEPD broth and incubated till mid-log phase and further incubated for 4 h in the presence of sub-inhibitory concentration of caspofungin. Cells were collected by centrifugation and vortexed with 1 mL of Trizol reagent with glass beads for 10 min with intermittent chilling. Then 0.2 ml of chloroform was added, and the upper layer was separated in a microcentrifuge tube after centrifugation at 13000 rpm for 15 min. An equal volume of chilled isopropanol was added to the cell lysate and precipitated at −80°C for 2 h. RNA was eluted in molecular grade water after centrifugation and quantified using Nanodrop Spectrophotometer (Thermo Scientific, Nanodrop 2000). First-strand cDNA was synthesized using the High Capacity cDNA kit (Thermofisher, Bengaluru, India). Quantitative real-time PCR was performed using the Power-up SYBR Green Master Mix (Applied Biosystems). Relative expression of each gene was calculated as fold-change in the presence of caspofungin calculated as 2^−^ΔΔCt (40)

**Table 3:**
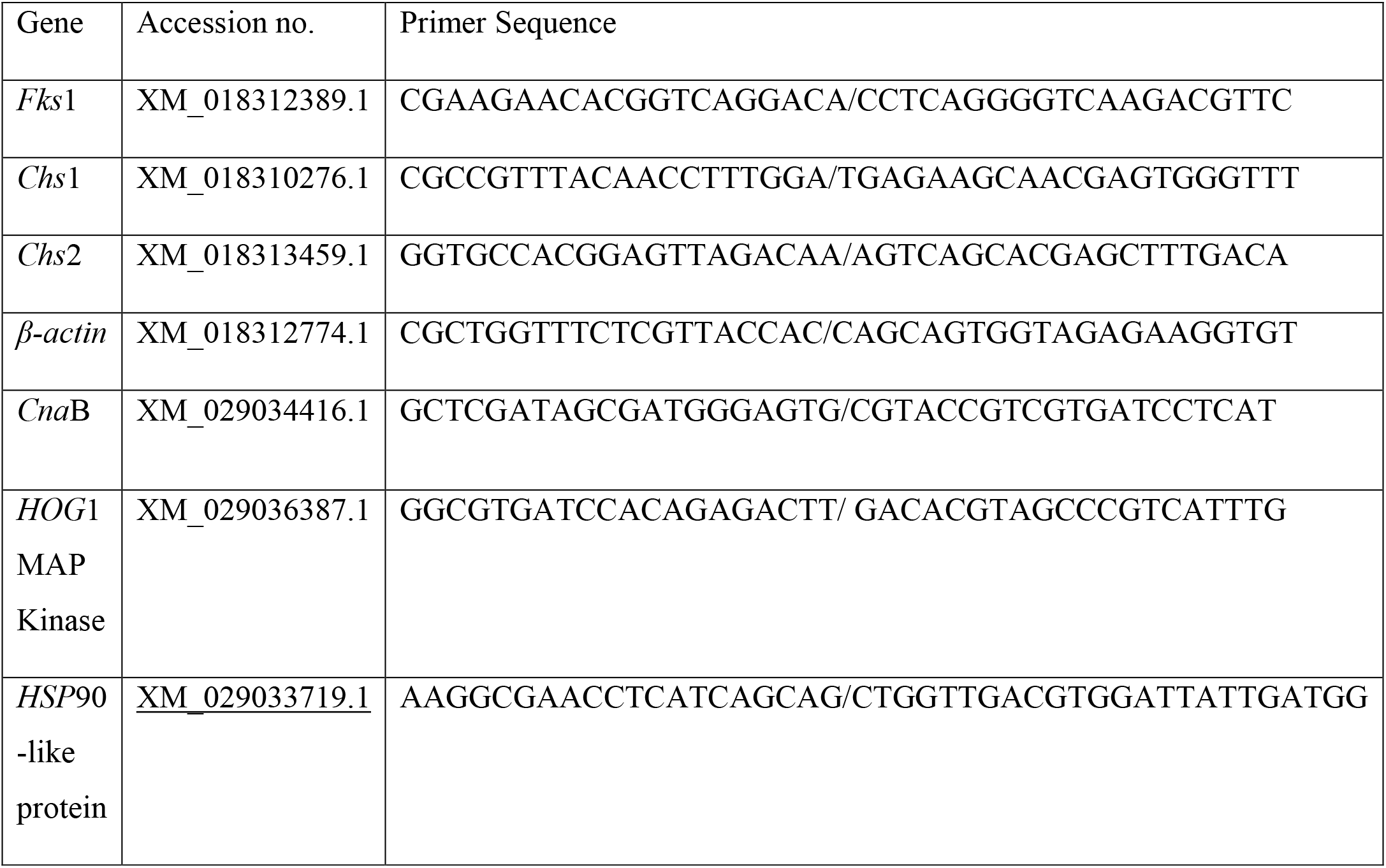
Primers used in the study

### Chitin content estimation

A previously described flowcytometry-based method was used to measure the baseline and caspofungin-inducible cell wall chitin contents (41). Cell suspensions in YEPD broth were exposed to respective sub-MIC caspofungin concentrations, as described above. Cells were washed twice with PBS, and approximately 1×10^6^ cells were treated with 2.5 mg/L calcofluor white (CFW) for 15 minutes. Cells were washed and subjected to flowcytometry using BD FACS Canto™ II (Becton Dickinson, San Jose, California, USA). Mean intensity (MI) of stained and unstained cells were measured, and the staining index was calculated using the following equation:

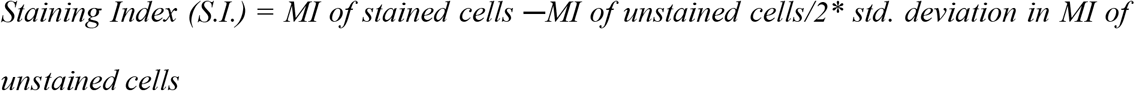

### Fluorescent Amplified Fragment Length Polymorphism

Amplified fragment analysis of six sequential isolates from a single patient was performed to assess the clonality of isolates as per our earlier described (8) method with a few modifications. Briefly, approximately 50 ng of genomic DNA was subjected to combined restriction-ligation reaction containing 50 pmol of *Eco*RI adapter, 50 pmol *Hind*III adapter, 2 U of *Hind*III (New England Biolabs, Beverly, MA, USA), 2 U of *Eco*RI (New England Biolabs) and one unit of T4 DNA Ligase in a total volume of 20 μL of 1X reaction buffer for one hour at 200 C. 1μL of the diluted restriction ligation mixture was used for amplification in a volume of 20 μL under the following conditions: 1 mM *Hind*III primer (5-Flu-GTAGACTGCGTACCCGTC-3), 1 mM *Eco*RI primer with one selective residues (5 - GATGAGTCCTGACTAA C-3), 0.2 mM of each dNTP and 1 U of Taq DNA polymerase (Roche Diagnostics) in 1X reaction buffer containing 1.5 mM MgCl2. Fragment analysis was done by capillary electrophoresis using LIZ-500 as internal size standard (Gene Scan 500, Applied Biosystems) and the typing data was analyzed by using BioNumerics software version 7.1 (Applied Maths, Ghent, Belgium). Phylogenetic tree was constructed by UPGMA (unweighted pair group method with arithmetic mean) with 1000 bootstrap replications in combination with Pearson’s correlation co-efficient.

### Growth Kinetics

Representative isolates with or without an *Fks*1 mutation were examined for any differential growth behaviour. Approximately, 1×10^6^ cells were inoculated in YEPD broth in a 96 well flat-bottom microtitre plate and incubated at 37 °C for 24 hrs with continuous shaking. Optical densities were recorded at 600 nm every hour in Epoch 2 Microplate Reader (Biotek, Agilent Technologies).

### Statistical analyses

The fold changes in the relative expression of *Chs*1, *Chs*2, and *Fks1* were analyzed by the non-parametric Kruskal-Wallis test. Post-hoc analysis was done using either Holm-Sidak’s or Dunn’s multiple comparison tests. The expression data are represented as median fold induction with interquartile range (Q1-Q3). The basal and inducible chitin contents expressed as staining index were analyzed using Wilcoxon matched-pairs signed-rank test. The Bland-Altman analysis was conducted to determine the congruence between fold changes in transcriptional levels of chitin synthase genes and staining index. Growth curves were analyzed by repeated measures two-way ANOVA with Turkey’s multiple comparisons test. The data were analyzed in GraphPad Prism version 6.0.

## Acknowledgment

We acknowledge Ujjwayini Ray, Apollo Gleneagles Hospitals, Kolkata, Dr Anjali Shetty, PD Hinduja National Hospital and Medical Research Centre, Hospital, Mumbai, Dr. Kamini Walia, ICMR, New Delhi and all those who sent *C. auris* isolates to the nodal centre as per the ICMR notification. We would also like to acknowledge technical support of Mr. Sourav Agnihotri. A part of this work was accepted to be presented in the 30^th^ meeting of European Congress on Clinical Microbiology and Infectious Diseases with the abstract number #9313.

All authors have no potential conflicts of interest. The funding agency did not participate or interfere in any stage of the study. All authors had full access to all trial data and take responsibility for the integrity of the data and the accuracy of the data analysis.

This work was supported by Indian Council of Medical Research (Grant # AMR/160/2018-ECD-II) New Delhi.

